# Time varying connectivity across the brain changes as a function of nicotine abstinence state

**DOI:** 10.1101/2020.01.25.917617

**Authors:** John R. Fedota, Thomas J. Ross, Juan Castillo, Michael R. McKenna, Allison L. Matous, Betty Jo Salmeron, Vinod Menon, Elliot A. Stein

**Affiliations:** Neuroimaging Research Branch, National Institute on Drug Abuse-Intramural Research Program, NIH, Baltimore, MD; Ohio State University, Columbus, OH; Geisel School of Medicine at Dartmouth, Hanover, NH; Stanford University School of Medicine, Stanford, CA

**Author notes:** equal contribution (John R. Fedota, Thomas J. Ross). Corresponding author: John R. Fedota & Elliot A. Stein, Neuroimaging Research Branch, National Institute on Drug Abuse-Intramural Research Program, NIH, 251 Bayview Blvd, Baltimore, MD 21224 (;).

## Abstract

**Background:** The Nicotine Withdrawal Syndrome (NWS) includes affective and cognitive disruptions whose incidence and severity vary across time during acute abstinence. However, most network-level neuroimaging employs static measures of resting state functional connectivity (rsFC), assuming time-invariance, and unable to capture dynamic brain-behavior relationships. Recent advances in rsFC signal processing allow characterization of “time varying connectivity” (TVC), which characterizes network communication between sub-networks that reconfigure over the course of data collection. As such, TVC may more fully describe network dysfunction related to the NWS.

**Methods:** To isolate alterations in the frequency and diversity of communication across network boundaries as a function of acute nicotine abstinence we scanned cigarette smokers in the nicotine sated and abstinent states and applied a previously-validated method to characterize TVC at a network and nodal level within the brain.

**Results:** During abstinence, we found brain wide decreases in the frequency of interactions between network nodes in different modular communities (i.e. temporal flexibility; TF). In addition, within a subset of the networks examined the variability of these interactions across community boundaries (i.e. spatiotemporal diversity; STD) also decreased. Finally, within two of these networks the decrease in STD was significantly related to NWS clinical symptoms.

**Conclusions:** Employing multiple measures of TVC in a within subjects’ design, we characterized a novel set of changes in network communication and link these changes to specific behavioral symptoms of the NWS. These reductions in TVC provide a meso-scale network description of the relative inflexibility of specific large-scale brain networks as a result of acute abstinence.

## Introduction

Acute nicotine abstinence is a key early hurdle to smoking cessation as most cessation attempts fail within a week of their target quit day^1^. These poor treatment outcomes are due, in part, to components of the Nicotine Withdrawal Syndrome (NWS) precipitated by acute smoking abstinence. The aversive symptoms of the NWS both dissuade smokers from attempting to quit^2^ and promote relapse via negative reinforcement—the relief of the withdrawal state^3–5^.

The clinical presentation of the NWS includes increases in negative affect^6,7^, lapses of attention^8–11^, and punctate craving for nicotine^6^. Critically, across both subjective and objective measures of NWS symptoms, variability in onset, time course, and phenomenology is consistently observed^12–17^. For example, craving increases early and again late in the day^15^, and negative affect spikes intermittently following cessation^17^. Symptoms are more variable in withdrawn smokers after quitting than before, suggesting that smoking may buffer or constrain aversive symptoms, which are then “unleashed” by cessation^18^. The incidence of these NWS symptoms increases with increased acute psychosocial stress^19–22^. The volatility in affective disruptions may also interact with the observed cognitive disruptions such that periodic lapses in attention—caused by subjective feelings of distress^23,24^ and nicotine craving^14^—drive abstinence-related cognitive decrements^8^.

As substance use disorder (SUD) is considered a brain circuit- and network-level disease^25,26^, the variability in the presentation of NWS clinical symptoms should be linked to dynamic changes in brain network communication and configuration over time. To this point, characterization of large-scale network communication in SUD populations generally show *increases* in resting state functional connectivity (rsFC) as a function of abstinence across a variety of circuits and brain networks^27,28^. However, considering the variability in clinical NWS symptom presentation, it is notable that these <u>static</u> rsFC studies have almost exclusively employed methods that assume unchanging network structure over time. This assumption limits the temporal resolution of the network communication described to the total data acquisition period and likely underspecifies the nature of the brain-based disruptions associated with the NWS.

Recent advances in fMRI time series signal processing allow characterization of “time varying connectivity” (TVC)^29–33^, which characterizes network communication between subnetworks that reconfigure over the course of data collection. Such characterizations of TVC are of particular relevance to the study of SUD, as ongoing, spontaneous (i.e. dynamic) brain activity has been related to the maintenance of brain circuit homeostasis^34,35^ via adaptive physiological response to external and internal perturbation. These homeostatic processes are impaired across a variety of SUDs, including nicotine dependence^36^. Specifically, these less adaptive compensatory responses to homeostatic challenges in SUD^37^ may be indexed by measures of TVC. Thus, we predict that the external perturbations of and stress induced by acute abstinence in SUD will be associated with reduced TVC.

To date most TVC studies have interrogated data from healthy individuals, and demonstrate reductions in TVC associated with poorer behavioral performance^31,38-40^ and increased negative affect^41^. Consistent with these affective and behavioral disruptions, a small but growing literature on SUD-related change in dynamic TVC has begun to appear^42–45^. These SUD studies consistently report a *reduction* in TVC associated with SUD, including a study from our lab reporting *decreases* in TVC as a function of acute nicotine abstinence^45^.

Here, we employ a TVC method developed and validated in healthy individuals^46^ allowing for characterization of TVC at both a brain network and nodal level. The observed changes in TVC are then related to changes in subjective and objective NWS symptoms in a within-subjects design. We hypothesize that acute nicotine abstinence will lead to decreased TVC in a set of brain networks and nodes previously associated with attentional processing and the NWS (Salience Network (SN), Frontoparietal Control Network (FPC), Default Mode Network (DMN))^26^ as well as those associated with emotional processing (insula-amygdala, anterior cingulate cortex, and ventral medial orbitofrontal cortex (vmOFC))^47^. Further, we hypothesize that decreased TVC will be associated with increased NWS symptom severity.

## Methods

### Participants

36 participants completed all experimental procedures. All participants were required to be right handed, between the ages of 18-60, free from active drug or alcohol abuse/dependence (other than current nicotine dependence), reporting no current psychiatric or neurological disorders, presenting no contraindications for magnetic resonance imaging (MRI) and able to abstain from alcohol and caffeine for 24 hours before each of two scan sessions. 11 of 36 participants were excluded from analysis for excessive head motion (average framewise displacement (FD) >0.2 mm^48^) in either of their two scans, resulting in data analyzed from n=25 participants (Table 1). Written informed consent was obtained in accordance with the National Institute on Drug Abuse (NIDA)-Intramural Research Program Institutional Review Board.

**Table 1:** Participant Demographics

|  | N=25 |
| --- | --- |
| Gender (M/F) | 15 / 10 |
| Age | $37.76 \pm 2.05$ |
| Race (AA/C/MR) | 11 / 13 / 1 |
| Education (years) | $13.2 \pm 0.41$ |
| IQ (WASI) | $103.56 \pm 2.62$ |
| FTND | $4.72 \pm 0.35$ |
| Cigarettes per day | $13.96 \pm 1$ |
| Age of Smoking Initiation | $15.88 \pm 0.76$ |
| Years smoked | $18.4 \pm 2$ |

*Values are n/n or mean ± SE.*
*AA=African American; C=Caucasian; MR=Multiple Races;*
*WASI= Wechsler Abbreviated Scale of Intelligence;*
*FTND=Fagerström Test of Nicotine Dependence*

### Experimental Design

The experiment followed a longitudinal, within-subjects design where each participant completed two MRI scanning sessions: one during *ad lib* sated smoking followed by an acute abstinence scan, with the last cigarette ~48 hours before scan. The order of the two scans was fixed, with sated preceding abstinence scan by an average of 60.5 days (median=22.1 days).

During both MRI scanning sessions participants completed an 8 minute, eyes open resting scan directly followed by a 25-minute version of the parametric flanker task (PFT). Data was acquired on a 3T Siemens Trio scanner (Erlangen, Germany) using a 12-channel head coil. All stimuli were presented using E-Prime software (Psychology Software Tools, Sharpsburg, PA), and behavioral responses were collected with a four button MRI-compatible button box (Fig 1).

**Figure 1:**
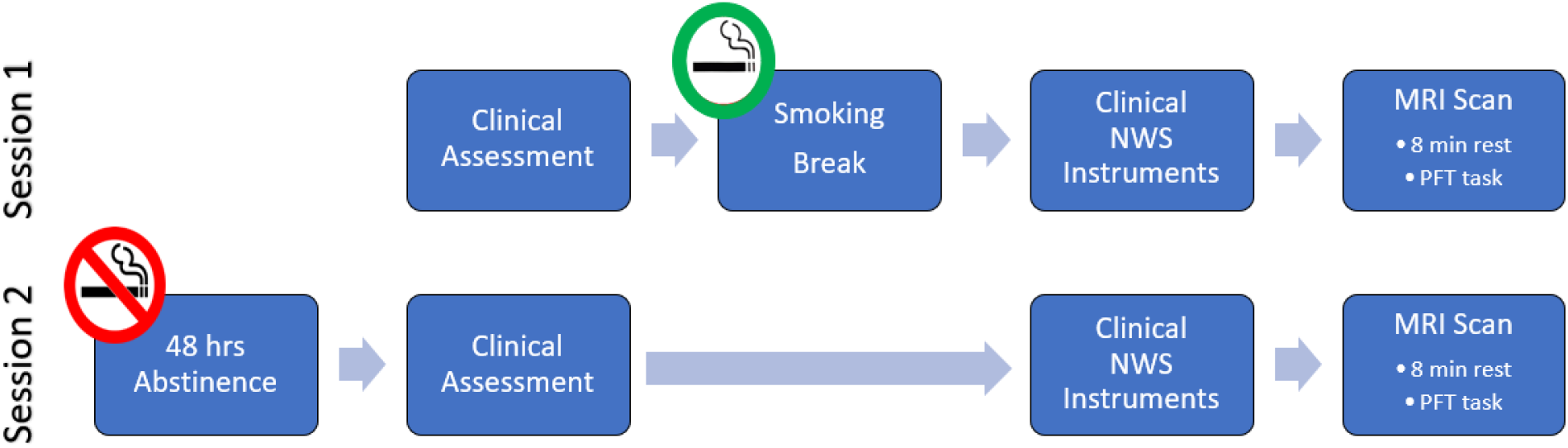
Longitudinal within-subjects design. Session 1=Sated smoking (last cigarette immediately prior to subjective clinical NWS instruments and ~45min prior to MRI scan). Session 2=acute nicotine abstinence (~48 hours of biologically-verified abstinence prior to subjective clinical NWS instruments and MRI scan).

### Clinical NWS Instruments Subjective ratings and analyses

Immediately prior to each scanning session, subjective ratings of withdrawal (Wisconsin Smoking Withdrawal Scale (WSWS))^49^, affect (Positive and Negative Affect Schedule (PANAS))^50^, perceived stress (Perceived Stress Scale (PSS))^51^, and craving (Tobacco Craving Questionnaire (TCQ))^52^ were assessed using previously-validated clinical instruments.

Subjective clinical instrument scores for each scan session were assessed. STATE (abstinence [–] sated) effects were calculated via paired t test for total score of WSWS, and PSS as well as factor scores for TCQ. Subscale scores for PANAS were submitted to a SUBSCALE*STATE repeated measures ANOVA.

### Behavioral performance and analyses

The PFT was used as an assay of attentional processing. The PFT is a modified version of the classic Eriksen flanker task designed to represent varying levels of demand for cognitive control driven by response conflict on a trial-by-trial basis. The procedural details of the task implementation in smokers have been described previously^53^. Behavioral performance on the PFT was quantified via correct reaction time (RT) and correct RT coefficient of variation (RTCV) (i.e. standard deviation of RT/mean RT). Counts of error type (Errors of Commission, Errors of Omission) were evaluated to assess selective attention as a function of nicotine abstinence. For errors of omission, STATE effects were quantified by paired t-test. For all other behavioral measures, values were submitted to a STATE (sated-abstinent) * DEMAND for cognitive control (high/medium/low) repeated measures ANOVA.

### MRI data acquisition and analyses

We followed the processing pipeline, including the analysis code, from the original TVC study^54^. (Fig 2) Raw data were minimally preprocessed using fmriprep (v0.4.5)^55^, with the first 10 frames discarded to account for scanner equilibrium. (See Supplemental Methods for full acquisition and preprocessing details)

**Figure 2:**
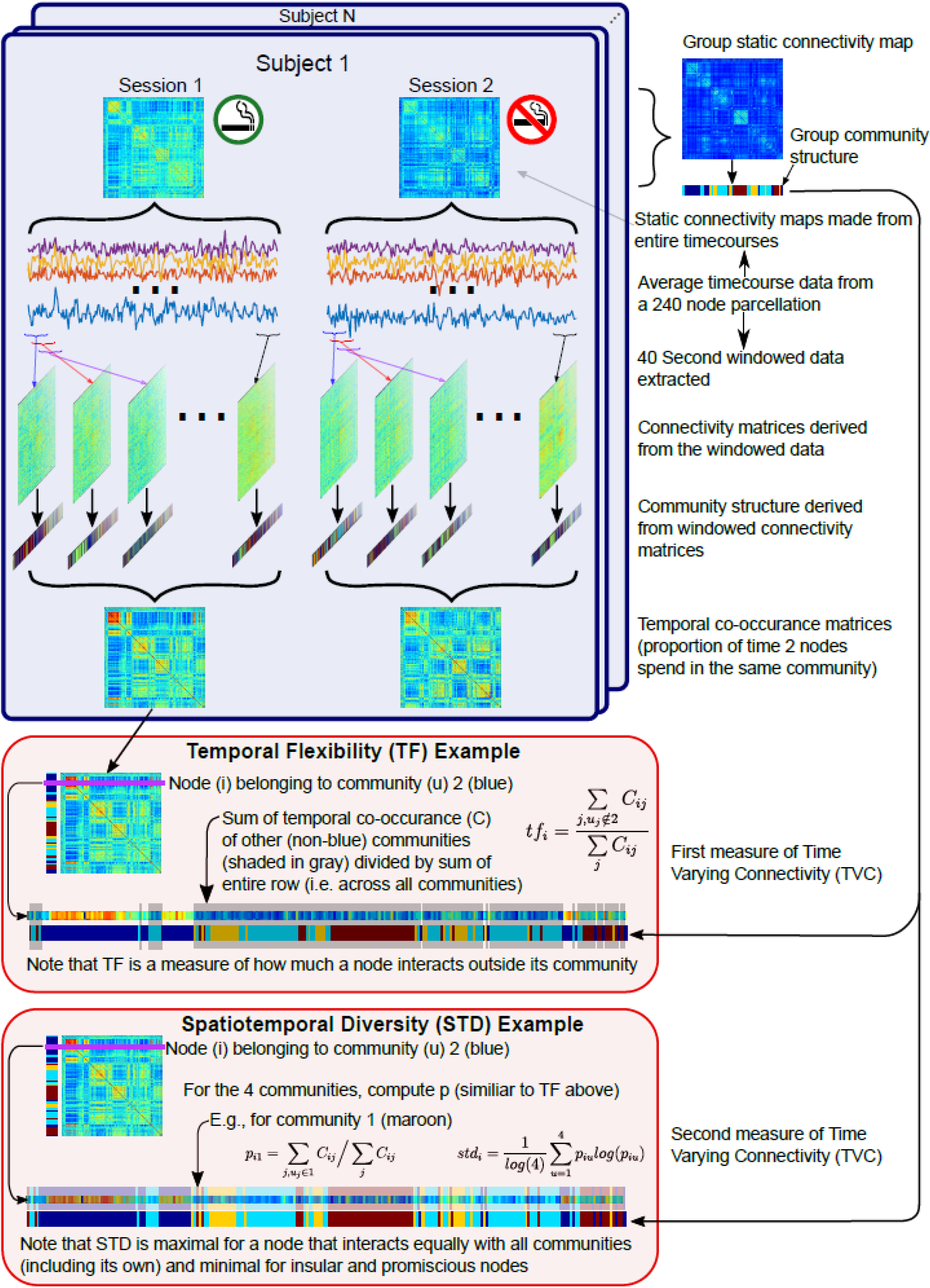
Time varying connectivity analysis pipeline. Average time courses were extracted from a 240 node parcellation (5mm radius spheres based on^63^ (see figure S1 for spatial distribution)). Static connectivity matrices from the entire time courses were combined to form a group connectivity matrix, and from this a community structure was derived using network theory. In our case, 4 communities were found, and nodal community membership is indicated by the colors in the bar adjacent to the group matrix. Separately, dynamic connectivity matrices were formed from 40 s windows of the time courses, their community structure calculated, and temporal co-occurrence matrices created consisting of the fraction of time that 2 nodes spend in the same community. These co-occurrence matrices were combined with the group community structure to form two measures of time-varying connectivity. The highlighted boxes give example calculations of these 2 measures for a hypothetical node belonging to community 2 (in this example colored blue in the group community structure).

The brain was parcellated into 264 nodes based on the canonical Power et al. scheme^59^, including supraordinate organization into 14 large-scale networks; 5mm spheres were placed at the center coordinates for each of the 264 ROIs. Based on the group EPI mask, any node with <50% voxel coverage was excluded, resulting in 240 nodes included in the current analysis. The 24 nodes from the original parcellation that are unused in the current study are in the anterior temporal lobe and OFC and belonged to the Default Mode and “Uncategorized” Networks (see Supplemental Methods and Fig S1 and Table S2 for excluded regions). For each scan, mean signals within the 240 nodes were extracted and high-pass filtered (f>.008 Hz) using a least-squares FIR filter (MATLAB function firls).

### Functional connectivity matrix (FCM)

At the individual subject and session levels, both a static and a dynamic FCM were created. Static FCM (240×240) were created using Pearson correlation on the entire 230-point timeseries and Fischer z-transformed. Dynamic FCM were created using a 40s sliding window, a 1 TR step, and exponentially decaying weights^60^. The weights were set to

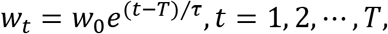

where *w_0_* is set such that the coefficients sum to one and τ is set to 1/3 of the window length. As with the static FCM, the weighed Pearson correlations were Fisher transformed. This resulted in 210 (240×240) dynamic FCM per individual and session.

### Community detection

Community detection was implemented using the Louvain algorithm implemented in the Brain Connectivity Toolbox (v2017_04_05)^61^, resulting in a community structure based on an optimal Q* parameter that maximizes intramodular connectivity while minimizing intermodular connectivity^62^. For each application, community detection was randomly initialized 100 times and the iteration with the largest Q* was retained. This community structure is thus data-driven and is not restricted by the *a priori* network membership imposed by the Power et al., parcellation^63^. Community detection was performed on group average and session average static FCM and, at the individual level, on the FCM from each of the 210 dynamic windows. The community structure derived from the group average static FCM was applied in the calculation of TVC metrics as described below.

### TVC metrics

Within each dynamic window an adjacency matrix was computed. In the adjacency matrix, if nodes *i* and *j* are in the same community, then cell (*i,j*) is 1 (or 0 if their communities differ). The adjacency matrices were averaged across the 210 dynamic windows to create a temporal co-occurrence matrix *C*. Thus, cell *C_ij_* quantified the proportion of time that nodes *i* and *j* spent in the same community, even as the overall community structure changed over time. Temporal co-occurrence matrices were computed for each individual in each session. The temporal cooccurrence matrix and group static FCM were then combined to derive two nodal-level measures of TVC as follows:

<u>Temporal flexibility (TF)</u> quantifies the degree to which a given node interacts outside of its reference community. For nodes *i* and *j*, participant *k* and session *l*, TF is:

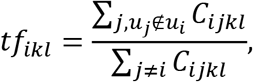

where *u_i_* is the community of node *i* from the group static FCM. Thus, TF represents the frequency of interactions a given node has outside of its static reference community divided by the total interactions a given node has with all other nodes.

<u>Spatiotemporal Diversity (STD)</u> quantifies the variability in a node’s interaction outside of its reference community. It is the normalized connection diversity^64^, using the temporal co-occurrence matrix instead of the FCM, and is inspired by Shannon entropy. For node *i*, participant *k* and session *l*, the STD is:

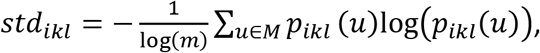

where *M* is the set of communities (numbering *m*), *p_ikl_*(*u*) = *s_ikl_, s_ikl_* is the sum of the temporal co-occurrence matrix for node *i*, participant *k* and session *l* and all communities and, similarly, *s_ikl_*(*u*) is the sum for community *u*. The log(*m*) term normalizes *std* to [0,1]. While TF and STD both measure similar constructs, they do differ. For example, a node can frequently interact with a single node outside its community (high TF, low STD) or interact frequently with a variety of nodes from other communities (high TF, high STD).

Network and Community values for TF and STD were calculated by averaging the nodal scores within each of the 14 *a priori* networks based on the Power et al. parcellation^63^ or the 4 detected communities (see Results) in the group static FC matrix, respectively to assess the robustness of the results to alternative network definitions.

### TVC session effects

Changes in TVC metrics (TF and STD) calculated at the network or community level as a function of STATE were assessed via repeated measures ANOVA with factors NETWORK and STATE (sated/abstinent) or COMMUNITY and STATE, with change in framewise displacement (FD) included as a covariate. Main effects of STATE or interactions (e.g. NETWORK*STATE) in the absence of an FD interaction (i.e. NETWORK*STATE*FD) were characterized with post-hoc comparisons, Bonferroni corrected for multiple comparisons at α<0.05.

### Correlation analyses

To relate changes in subjective withdrawal symptoms and/or behavior to changes in TVC (at the network, community, or node levels), statistical models were built via robust linear regression using the lmrob function from R package *robustbase*^65^. Based on the variability inherent in subjective reports, robust regression was employed to provide statistical estimates less sensitive to the presence of outliers in the data. As with the TVC session effects, change in FD was included in the regression model (Δ behavior ~ Δ brain + ΔFD) to control for relationships confounded by residual changes in head motion across scan sessions.

## RESULTS

### Abstinence and clinical NWS assessments

#### *Physiological* assessment

Consistent with self-reports of ~48 hours of full nicotine abstinence (50.2±9.7 hrs), expired CO was significantly reduced during abstinence (sated scan (22.8±1.6) ppm; abstinent scan (2.5±0.2) ppm; F(1,24)=186.8 p<.0001).

#### Clinical Instruments

Subjective ratings of stress, affect and withdrawal were all modulated by acute nicotine abstinence; subjective ratings of craving, however, were not. PSS showed a STATE effect (F(1,24)=4.50, p<.05) such that perceived stress was greater during abstinence. PANAS showed a STATE*SUBSCALE (positive/negative affect) interaction (F(1,24)=18.92, p< .001). Follow-up tests showed positive affect decreased during abstinence (F(1,24)=21.02, p<.0001) while negative affect was unchanged (F(1,24)=2.42, p=.13). WSWS total score showed a strong trend level STATE effect (F(1,24)=4.10, p=.054) with increased ratings of withdrawal during abstinence. None of the four TCQ factors showed a STATE effect (all F’s < 0.95) (Figs 3A, S2)

**Figure 3:**
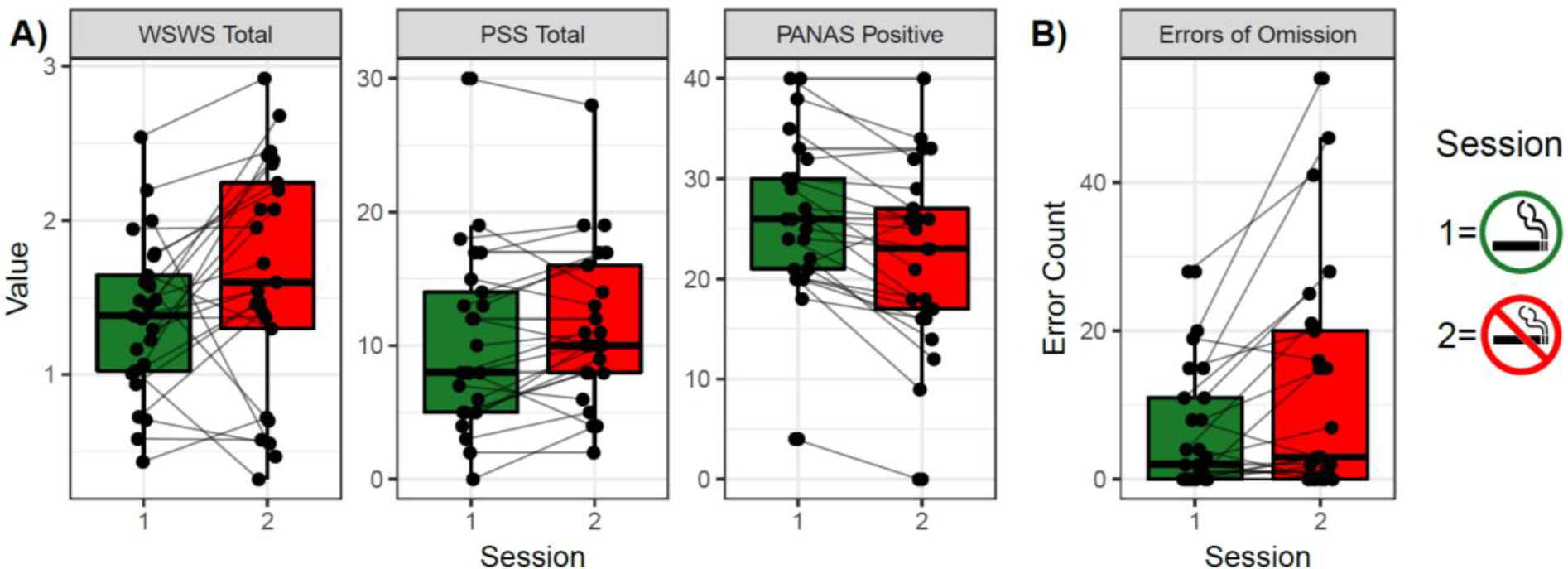
Abstinence induced change in clinical measures of the Nicotine Withdrawal Syndrome. A) subjective (WSWS, PSS, PANAS Positive) and B) behavioral (Errors of Omission) STATE ([abstinence-sated]) effects. See Supplemental Materials for all clinical measure results. Dots represent individual subject data in each session. WSWS=Wisconsin Smoking Withdrawal Scale; PSS= Perceived Stress Scale; PANAS=Positive and Negative Affect Schedule; Session 1=smoking satiety, Session 2= smoking abstinence

#### Behavioral performance

In the Parametric Flanker Task (PFT), Errors of Omission, showed a main effect of STATE such that omissions increased during abstinence (sated scan (6.08±7.88); abstinent scan (12.1±15.9); (F(1,24)=8.62, p <.01). No other measure of behavioral performance (RT, RTCV, Errors of Commission) showed an effect of STATE (all F’s < 2.20) or a STATE*DEMAND interaction (all F’s < 0.94) (Figs 3B, S3)

### Neuroimaging results

Even with the removal of 11 subjects due to excessive movement (FD> 0.2 mm), residual FD showed an effect of STATE (t(24)=3.79, p<.001) such that average FD was greater during the abstinence (0.14 mm) than the sated (0.11 mm) scan. Although these FD levels fall well below the floor effect previously described^48^, change in FD was added as a covariate to all subsequent neuroimaging analyses of STATE effects.

#### Temporal Flexibility (TF) and Spatiotemporal Diversity (STD): a priori network analyses

When nodal TVC values were averaged across the *a priori* networks (Power et al.^63^), *TF* showed a main effect of STATE (F(1, 23)= 8.81, p<.01) such that TF was reduced in abstinence vs. sated condition, while *STD* showed a STATE * NETWORK interaction (F(3.71, 85.24)=2.60, p<.05). Subsequent post hoc tests, corrected for multiple comparisons across the 14 networks, showed reductions in STD for the Cingulo-opercular Control network (COC) (F(1,23)=17.45, p<.005), default mode network (DMN) (F(1,23)=16.84, p<.01) and the “uncategorized” network (UNC) (F(1,23)=14.58, p<.05); there was a strong trend level effect for the Salience Network (SN) (F(1,23)= 9.88, p=.05. In each case, network STD was reduced during abstinence vs. sated (Fig 4, S4, S5).

**Figure 4:**
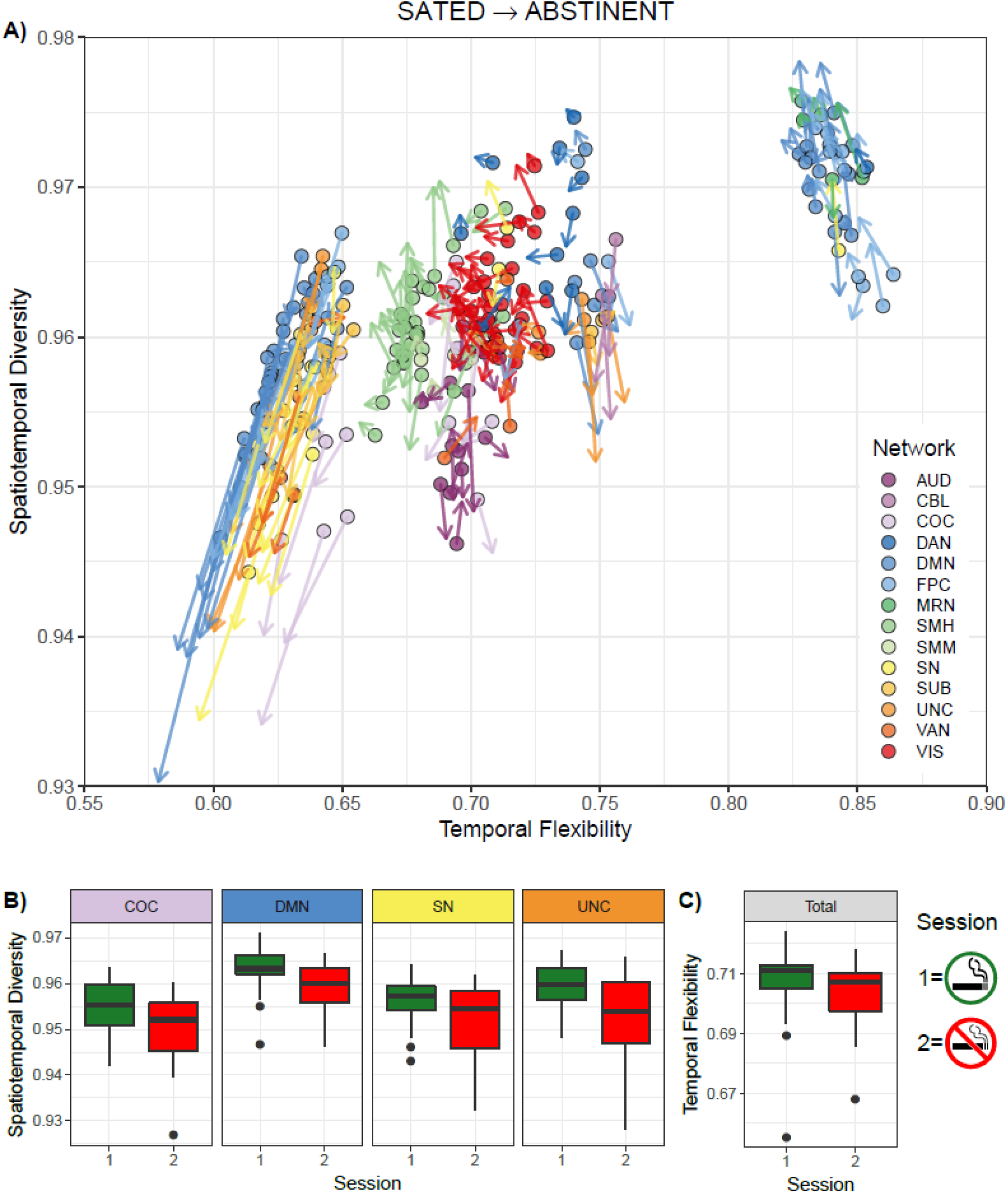
Time varying connectivity change as a function of abstinence. A) Nodal results. Each circle represents a single node (240 total) from the whole brain parcellation. Color of the circle indicates <u>a priori</u> network membership (56). Vectors indicate the magnitude of change in Temporal Flexibility (TF) and spatiotemporal diversity (STD) as a function of nicotine abstinence. B) Network results. Average STD value across nodes constituting each of the four a priori networks showing a decrease as a function of abstinence. C) Whole brain results. Average TF value across all 14 networks decreased as a function of abstinence. Network abbreviations: AUD=Auditory; CBL= Cerebellar; COC=Cingulo-Opercular Control; DAN=Dorsal Attention; DMN=Default Mode; FPC=Fronto-Parietal Control; MRN=Memory Retrieval; SMH=Somatomotor Hand; SMM=Somatomotor Mouth; SN= Salience; SUB=Subcortical; UNC=Uncategorized; VAN=Ventral Attention; VIS=Visual

#### TF and STD: data driven reference community analyses

When nodal TVC values were averaged across detected community membership, TF showed a main effect of STATE (F(1,23)=12.92, p<.005) such that TF was reduced during abstinence. Additionally, STD showed a STATE*COMMUNITY interaction (F(2.24, 51.54)=6.73, p<.005) such that only one of the four identified communities showed a reduction in STD as a simple main effect of STATE (F(1,23)=18.00, p<.005) (Fig S6).

### Correlations between behavioral measures and clinical reports with neuroimaging data

#### Subjective measures of withdrawal: Perceived Stress Scale (PSS)

Of the networks that showed a STATE effect (i.e. [abstinence (-) satiety]) on STD, change in UNC was positively related with change in PSS (p<.004) (Fig 5A).

**Figure 5:**
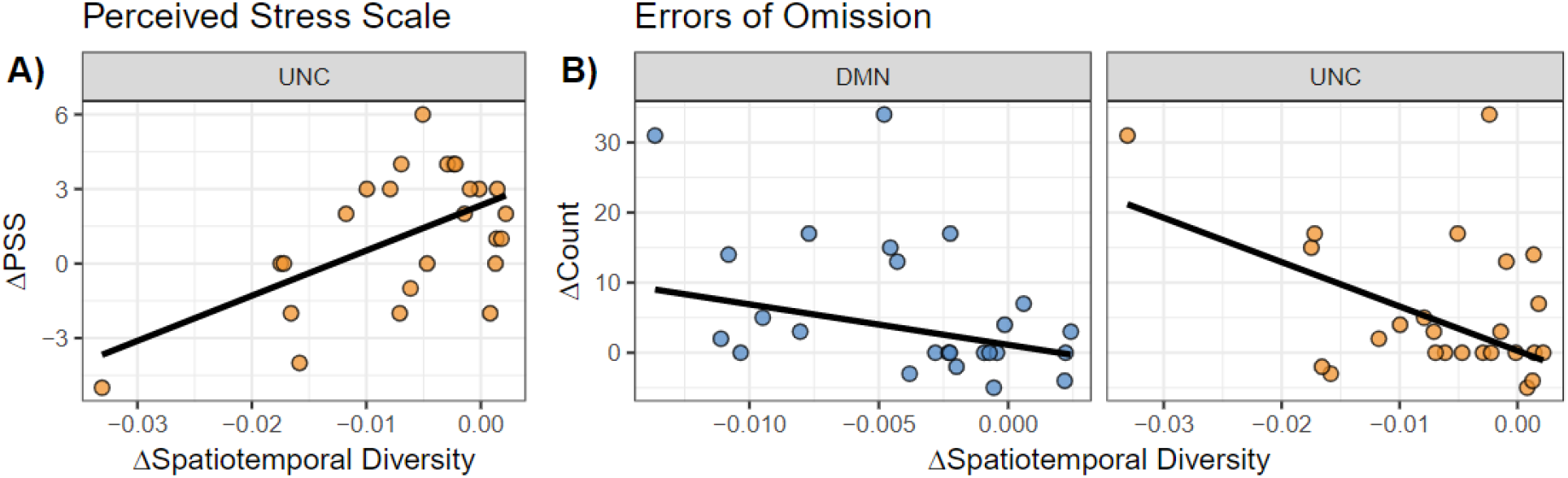
Correlations between abstinence-induced changes ([abstinent-sated]) in Spatiotemporal Diversity (STD) and clinical Nicotine Withdrawal Syndrome symptoms. A) Relationship with subjective report. A decrease in STD in the “Uncategorized” network is significantly related to a decrease in Perceived Stress Scale. B) Relationship with behavior. Decreases in STD in the Default Mode and “Uncategorized” networks are significantly related to an increase in errors of omission in the Parametric Flanker Task. DMN=Default Mode; UNC= Uncategorized; PSS=Perceived Stress Scale

#### Behavioral measures of withdrawal: Errors of Omission

Of the networks that showed a STATE effect on STD, change in STD within the DMN and UNC were negatively related with change in Errors of Omission (DMN, p<.02; UNC, p<.03) (Fig 5B).

## Discussion

We examined alterations in whole brain resting state TVC as a function of acute nicotine abstinence vs. satiety. We identified reductions in the frequency and variability of interactions between nodes within larger networks across the brain that point to an abstinence-precipitated allostatic alteration in brain dynamics. Brain wide *decreases* in the frequency of interactions between network nodes in different modular communities (i.e. TF) were observed following 48 hours of smoking abstinence. In addition, within a subset of the 14 *a priori* large-scale networks examined—COC, DMN, SN, UNC—the variability of these interactions (i.e. STD) across community boundaries was also *decreased*. Finally, within two of these networks (DMN and UNC), the decrease in STD was significantly related to NWS clinical symptoms. Thus, reductions in TVC during early abstinence, when most treatment failures are seen, appear to characterize a systems-level dysfunction that may be a key mechanistic component of the NWS.

### Temporal Flexibility (TF)

The frequency of a given network’s interaction with nodes outside of its reference community, defined as TF, demonstrated a brain-wide reduction in inter-community participation across both *a priori* networks and *data driven* community parcellations during abstinence. This suggests a maladaptive increase in segregation of individual networks and commensurate reductions in communication efficiency across the brain regardless of the network categorization employed. In healthy populations, such increased network segregation is associated with a) specialized processing and a lack of adaptability^66–68^, b) is seen during performance of less demanding or automatic tasks^69–71^ and c) promotes learned associations at the expense of flexible exploratory behavior^38,67,72,73^. In the case of SUD, acute abstinence enhances such segregation by reducing the frequency of communication across subnetwork boundaries.

This observed reduction in TF during abstinence can be interpreted within the allostatic overload framework^74,75^. During allostatic overload, repeated exposure to stress precipitates a dysregulated physiological response in an effort to restore homeostatic stability. However, this maladaptive response reduces stability and flexibility of the regulatory response^37,76^. As TVC is a key modulator of metastability within the complex dynamic whole brain network^38,77,78^, a reduction in the flexibility of connections over time is consistent with allostatic load. That is, the stress of acute abstinence is associated with a pervasive, brain-wide decrease in periodic communication between otherwise segregated network communities.

A putative mechanism for the observed decrease in TF is a reduction in synaptic plasticity during acute nicotine abstinence. Normally functioning synapses use metaplasticity to titrate levels of long-term potentiation and depression within a homeostatic dynamic range, thus avoiding run away strengthening or weakening of connections^79^. In contrast, SUD is associated with reduced synaptic plasticity, reduced dynamic response to stressors, and more rigid synaptic connections^80,81^. Thus, in SUD—especially during withdrawal—the brain is in a potentiated state and appears unable to reconfigure in response to environmental demands. Related clinical evidence illustrates an inability to induce synaptic plasticity in motor circuits via non-invasive brain stimulation during nicotine abstinence^82,83^. While these prior results describe reductions in plasticity of specific circuits, the current findings provide a meso-scale description of a similar phenomenon.

### Spatiotemporal Diversity (STD)

In contrast to the broad reduction in TF, a focused reduction in STD—the variability of interactions between a given set of nodes and the rest of the brain^46,84^—was observed in a subset of the networks or communities tested. Only four networks (COC, DMN, SN, and UNC) displayed a limited repertoire of interactions with other brain areas, and these interactions were less frequent during abstinence. Three of the 4 networks that display reduced STD during abstinence have been previously implicated in cognitive control (SN and COC)^85,86^, tonic alertness (COC and DMN)^87,88^ and abstinence-related dysfunction in SUD (SN and DMN)^26,89,90^. Further, the observed focal decreases in STD coincide with the sites of highest nicotinic acetylcholine receptor (nAchR) expression in the human brain (COC and SN)^91^, implicating disrupted nicotinic signaling dynamics in the reduction of STD during abstinence.

Beyond an overlap with the expression of nAchR’s, the networks displaying reduced STD during abstinence are canonical members of the “rich club” architecture of the brain^92^. The rich club consists of a group of densely interconnected nodes that have been shown to coordinate the integration of information processed locally throughout the brain^93–95^ and provide a stable scaffold upon which dynamic reconfigurations in the brain’s network structure in response to task demands or arousal occur^67,77,78,96^. Indeed, hubs within the rich club have been identified in the SN, COC, and DMN^92,94,95^; additional nodes in the orbitofrontal cortex (OFC; part of the UNC network in the current study) are part of the dynamical workspace of binding nodes^97^ that serve to integrate disparate processing across the brain as a compliment to the rich club.

During acute abstinence the integration of locally processed information across the whole brain network is disrupted as the networks associated with the rich club do not interact as broadly with other nodes across the brain as indexed by the reduction in STD for the four networks identified. The co-occurrence of reductions in the frequency (TF) and variability (STD) of communication in only these networks may have an outsized impact in promoting segregation at the cost of integration during abstinence, leading to the affective and attentional disruptions of NWS.

Strengthening this interpretation and the importance of hub regions displaying varied interactions with the rest of the brain, a subset of these observed reductions in STD was directly correlated with the clinical presentation of the NWS. Decreases in STD within both the DMN and UNC networks were negatively correlated with Errors of Omission in an attentional control task. That is, the greater the decrease in STD in DMN or UNC, the larger the increase in the number of Errors of Omission. These findings are consistent with disruptions in vigilance reported during abstinence^8,11^. Further, in healthy populations, optimal task performance is associated with integration across otherwise segregated networks in the brain^98^ and sparse connectivity across networks/communities has been previously related to self-reported fatigue^99^. Thus, the lapses in attention seen in abstinence and related to decreased communication across the brain fits with these normative findings.

In contrast to the intuitive relationship between decreased STD and increased attentional disruption, the relationship between decreased STD and affective symptoms of NWS is less clear. The positive correlation between abstinence induced changes within UNC network STD and perceived stress shows that as STD decreases, perceived stress also decreases. Our original hypothesis predicted increased NWS symptoms with decreased TVC. However, SUD is strongly associated with impairments in interoceptive processing^100–103^, including decreased activation to interoceptive cues in the OFC—a constituent node of the UNC network^104^. These findings plus the well-characterized dissociation between physiological arousal and self-report in smokers^13,105^ (i.e. Nesbitt’s Paradox^106^) suggest that the subjective point estimates of NWS employed in the current study may have been suboptimal in characterizing changes in the intensity (not to mention the frequency) of affective disruptions associated with changes in TVC. Future studies should consider using longitudinal measures of subjective NWS—for example, employing ecologic momentary assessment (EMA) methods^15,16,107,108^— to better characterize the volatility of these subjective symptoms during abstinence and relate them to the described decreases in TVC. To this point, recent evidence in depressed participants described decreased SN flexibility associated with increased affective disruption as measured via asynchronous ambulatory assessments^109^.

Taken together, the current study describes both broad and focused reductions in TVC during acute nicotine abstinence. These decreases are in contrast to the broad increase in static rsFC previously reported^27,28^, and identify specific networks whose function is impaired by abstinence. Further, these reductions in TVC create a meso-scale network description, linking prior evidence of reductions in synaptic plasticity with large-scale theories of allostatic load and relative inflexibility of network response to the acute stress of abstinence. By employing multiple measures of TVC in a within subjects’ design, we characterize a novel description of changes in network communication and link these changes to specific behavioral symptoms of the NWS. Moving forward, treatments focused on reversing or mitigating the effects of the observed network stasis on attentional processes during abstinence may be of interest.

## Acknowledgements

We thank Kim Slater, Bridget Moynihan, and Kevin Noemer for conducting the smoking cessation treatment sessions. We thank Tianwen Chen and Xiaoyu Ding for assistance with data analysis scripts.

Supported by the Intramural Research Program of the NIDA/NIH, FDA grant NDA13001-001-00000 to EAS, and NIH grant NS086085 to VM.

The authors report no biomedical financial interests or potential conflicts of interest

