## Supplemental Materials (methods, figures, tables) for "Time varying connectivity across the brain changes as a function of nicotine abstinence state"

### Supplemental Material

### Experimental Procedures

Upon arrival for each MR scanning session, participants underwent a brief clinical assessment including a urine test for recent use of illicit drugs (opiates, oxycodone, benzodiazepines, buprenorphine, cocaine, amphetamines/methamphetamines, THC, methadone, PCP, and MDMA) and Breathalyzer test for recent alcohol use. Positive tests were exclusionary for all drugs, except THC; positive urine tests for THC were followed by the Drug Evaluation and Classification neuromotor exam to determine if the participant was acutely intoxicated; a positive neuromotor exam was exclusionary. For the abstinent scan, compliance was assessed via both self-report and expired CO with ≥ 4ppm indicating a failure to maintain nicotine abstinence and need to reschedule the abstinence scan.

Following clinical assessments and immediately before administration of clinical instruments, to control for plasma nicotine levels during the sated scan, all participants were given a <10-minute smoking break, which resulted in last cigarette smoked an average of 42.7 minutes (SD=8.0) before MRI scanning.  This smoking break was omitted prior to the abstinent scan.

### Neuroimaging Procedures

#### *Data Acquisition*

During the 8-minute resting state scan, a T2*-weighted, single-shot gradient echo, echo planar imaging sequence sensitive to blood oxygenation level-dependent (BOLD) effects was used to acquire whole brain data on a 3T Siemens Trio scanner (Erlangen, Germany) using a 12-channel head coil.  Oblique axial slices (39, 4mm thick slices with no gap; 30° to anterior commissure-posterior commissure line) were acquired (240 volumes; repetition time (TR)=2000ms; echo time (TE)=27ms; flip angle (FA)= 78°; field of view 220 x 220 $mm^2$; image matrix 64 x 64). High resolution oblique-axial T1-weighted structural images were also acquired using a 3D magnetization-prepared rapid gradient-echo (MPRAGE) sequence (TR=1900ms; TE 3.51ms; TI=900ms; FA=9°; voxel size=1 $mm^3$).

*Fmriprep methods*

*(text in this section auto generated via https://fmriprep.readthedocs.io/en/stable/citing.html)*

Results included in this manuscript come from preprocessing performed using FMRIPREP version 0.4.5 [1, 2, RRID:SCR_016216], a Nipype [3, 4, RRID:SCR_002502] based tool. Each T1w (T1-weighted) volume was corrected for INU (intensity non-uniformity) using `N4BiasFieldCorrection` v2.1.0 [5] and skull-stripped using `antsBrainExtraction.sh` v2.1.0 (using the OASIS template). Spatial normalization to the ICBM 152 Nonlinear Asymmetrical template version 2009c [7, RRID:SCR_008796] was performed through nonlinear registration with the `antsRegistration` tool of ANTs v2.1.0 [8, RRID:SCR_004757], using brain-extracted versions of both T1w volume and template. Brain tissue segmentation of cerebrospinal fluid (CSF), white-matter (WM) and gray-matter (GM) was performed on the brain-extracted T1w using `fast` [17] (FSL v5.0.9, RRID:SCR_002823).

Functional data was slice time corrected using `3dTshift` from AFNI v16.2.07 [11, RRID:SCR_005927] and motion corrected using `mcflirt` (FSL v5.0.9 [9]). This was followed by co-registration to the corresponding T1w using boundary-based registration [16] with six degrees of freedom, using `flirt` (FSL). Motion correcting transformations, BOLD-to-T1w transformation and T1w-to-template (MNI) warp were concatenated and applied in a single step using `antsApplyTransforms` (ANTs v2.1.0) using Lanczos interpolation.

Physiological noise regressors were extracted applying CompCor [18]. Principal components were estimated for the two CompCor variants: temporal (tCompCor) and anatomical (aCompCor). A mask to exclude signal with cortical origin was obtained by eroding the brain mask, ensuring it only contained subcortical structures. Six tCompCor components were then calculated including only the top 5% variable voxels within that subcortical mask. For aCompCor, six components were calculated within the intersection of the subcortical mask and the union of CSF and WM masks calculated in T1w space, after their projection to the native space of each functional run. Frame-wise displacement [19] was calculated for each functional run using the implementation of Nipype.

Many internal operations of FMRIPREP use Nilearn [22, RRID:SCR_001362], principally within the BOLD-processing workflow. For more details of the pipeline see https://fmriprep.readthedocs.io/en/latest/workflows.html

*Additional preprocessing*

Differences in head motion are a significant confounding factor in MRI analysis [23,24], including in TVC [25], although the effects of motion in TVC may be limited [26]. Thus, in addition to removing subjects with mean FD>0.2 mm, to further control for residual head motion, 6 motion parameters and their temporal derivatives were regressed from the data. Data was then spatially smoothed with a 6mm FWHM Gaussian kernel. To address non-specific signal confounds, identified WM and CSF time-series derived from a 30-component ICA analysis using Melodic (FSL, version 3.14) were regressed out. Importantly, the average change in FD between the sated and abstinent scan was included in all STATE models as a covariate.

Supplemental Figures

Figure S1: *Brain parcellation employed*

Figure S2: *All subjective clinical instruments assessed*

Figure S3: *All behavioral clinical instruments assessed*

Figure S4: *Averaged time varying connectivity results*

Figure S5: Nodes within each network showing state effect on time varying connectivity

Figure S6: *Time varying connectivity change as a function of abstinence displayed by group community segmentation*

Figure S1: *Brain parcellation employed*

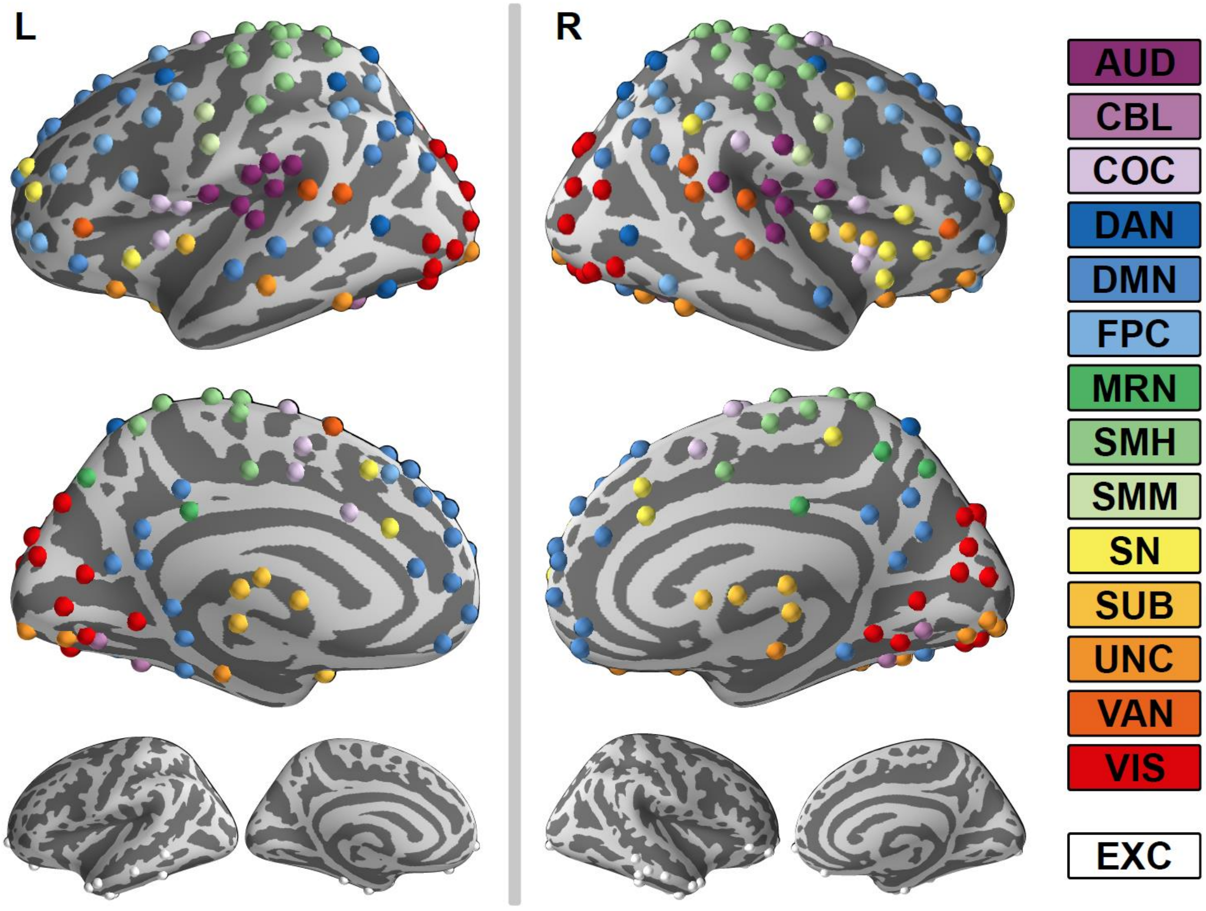


*Figure S1: Brain parcellation employed. N=240, 5mm spheres placed throughout the brain and supraordinate large scale a priori network organization denoted by color. White spheres (small surfaces, n=24) indicate nodes from original Power et al. 2011 parcellation that were excluded from the current study due to insufficient voxel coverage in field of view. See also supplemental tables 1 and 2.*

*AUD=Auditory; CBL= Cerebellar; COC=Cingulo-Opercular Control; DAN=Dorsal Attention; DMN=Default Mode; FPC=Fronto-Parietal Control; MRN=Memory Retrieval; SMH=Somatomotor Hand; SMM=Somatomotor Mouth; SN= Salience; SUB=Subcortical; UNC=Uncategorized; VAN=Ventral Attention; VIS=Visual; EXC=Excluded*

Figure S2: *All subjective clinical instruments assessed*

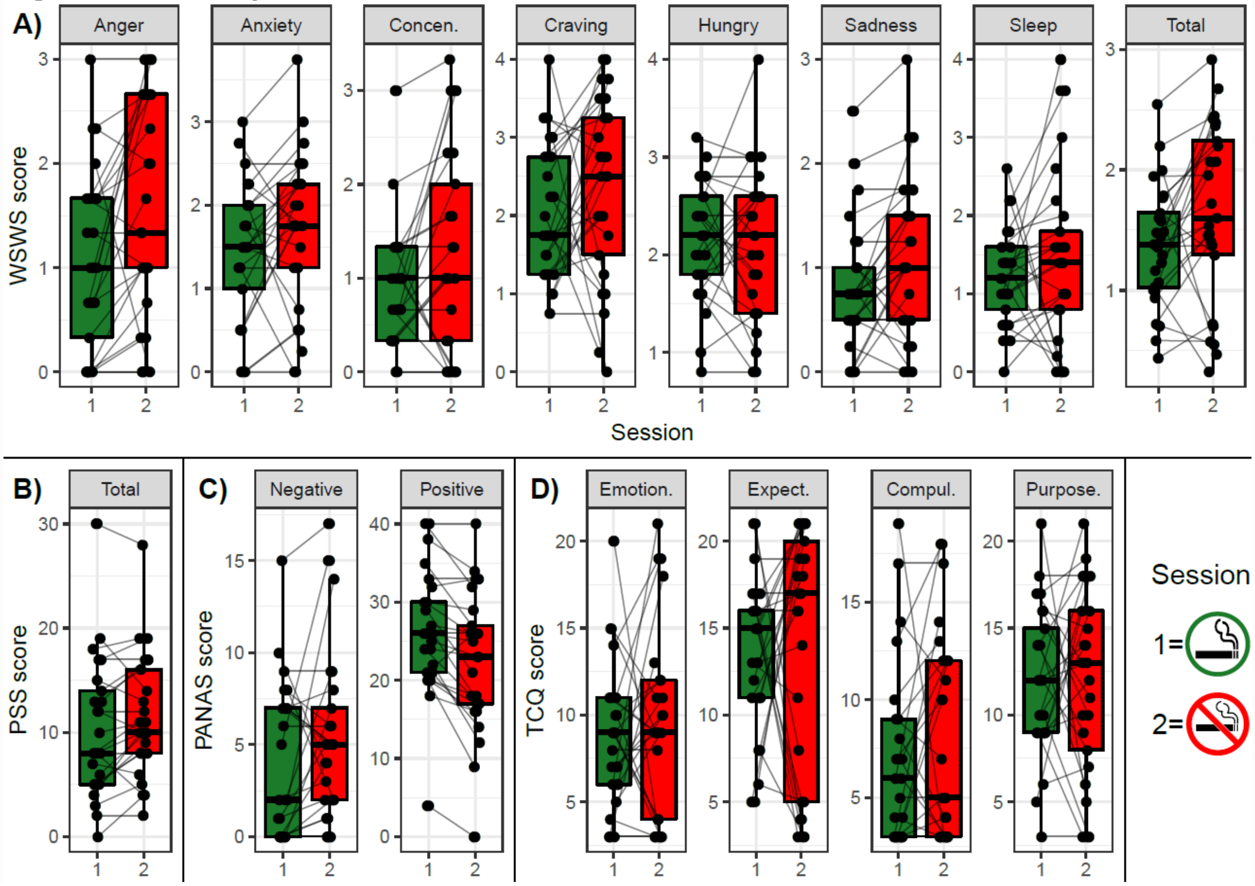


*Figure S2: All subjective clinical instruments assessed. A) Wisconsin Smoking Withdrawal Scale (WSWS) B) Perceived Stress Scale (PSS) C) Positive and Negative Affect Schedule (PANAS) D) Tobacco Craving Questionnaire (TCQ).*

Figure S3: *All behavioral clinical instruments assessed*

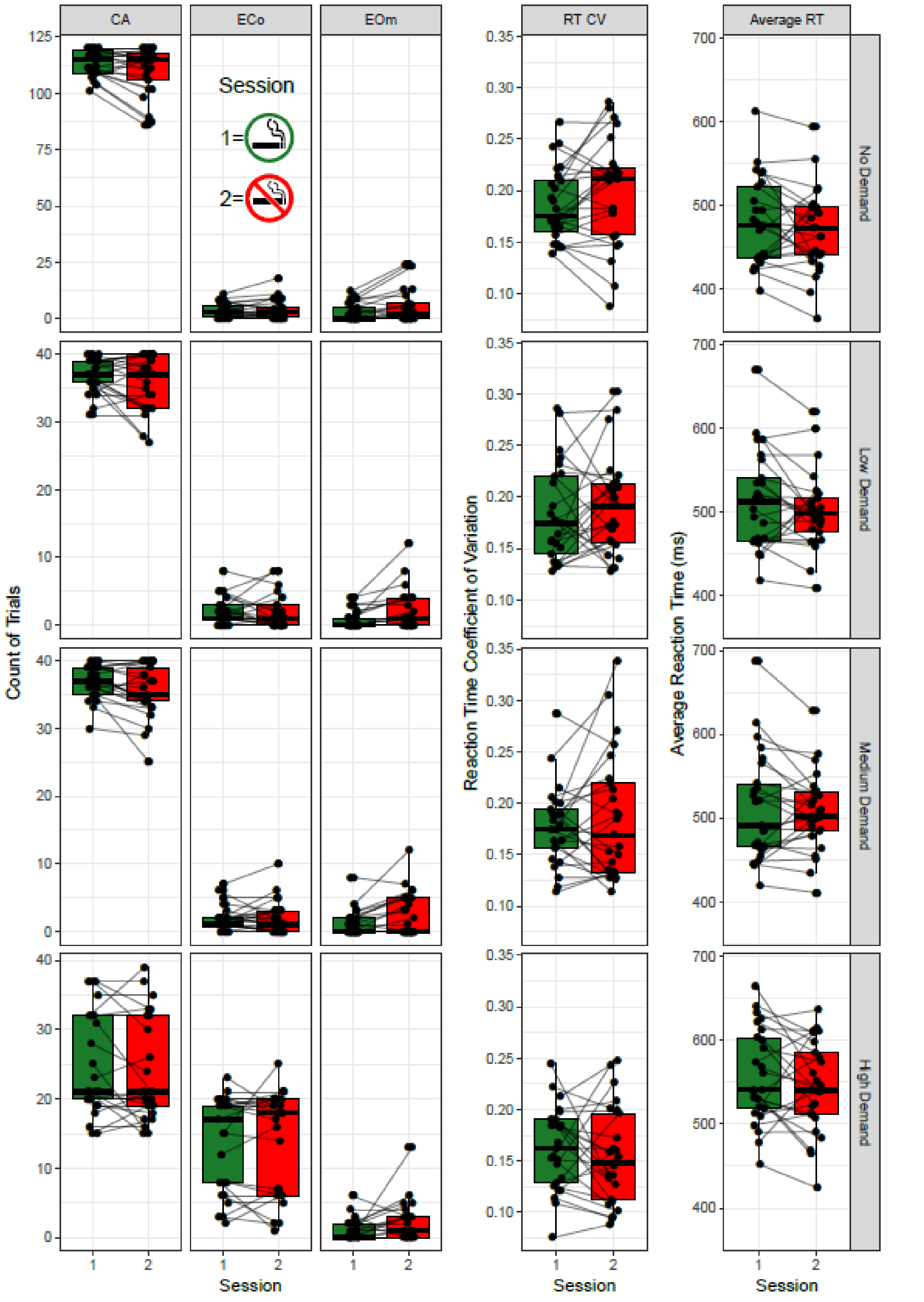

*Figure S3: All behavioral clinical instruments assessed.*

*CA=correct answers; ECo= errors of commission; EOm= errors of omission; RT= reaction time; RT CV= reaction time coefficient of variation (standard deviation of RT/mean RT)*

Figure S4: Averaged time varying connectivity results

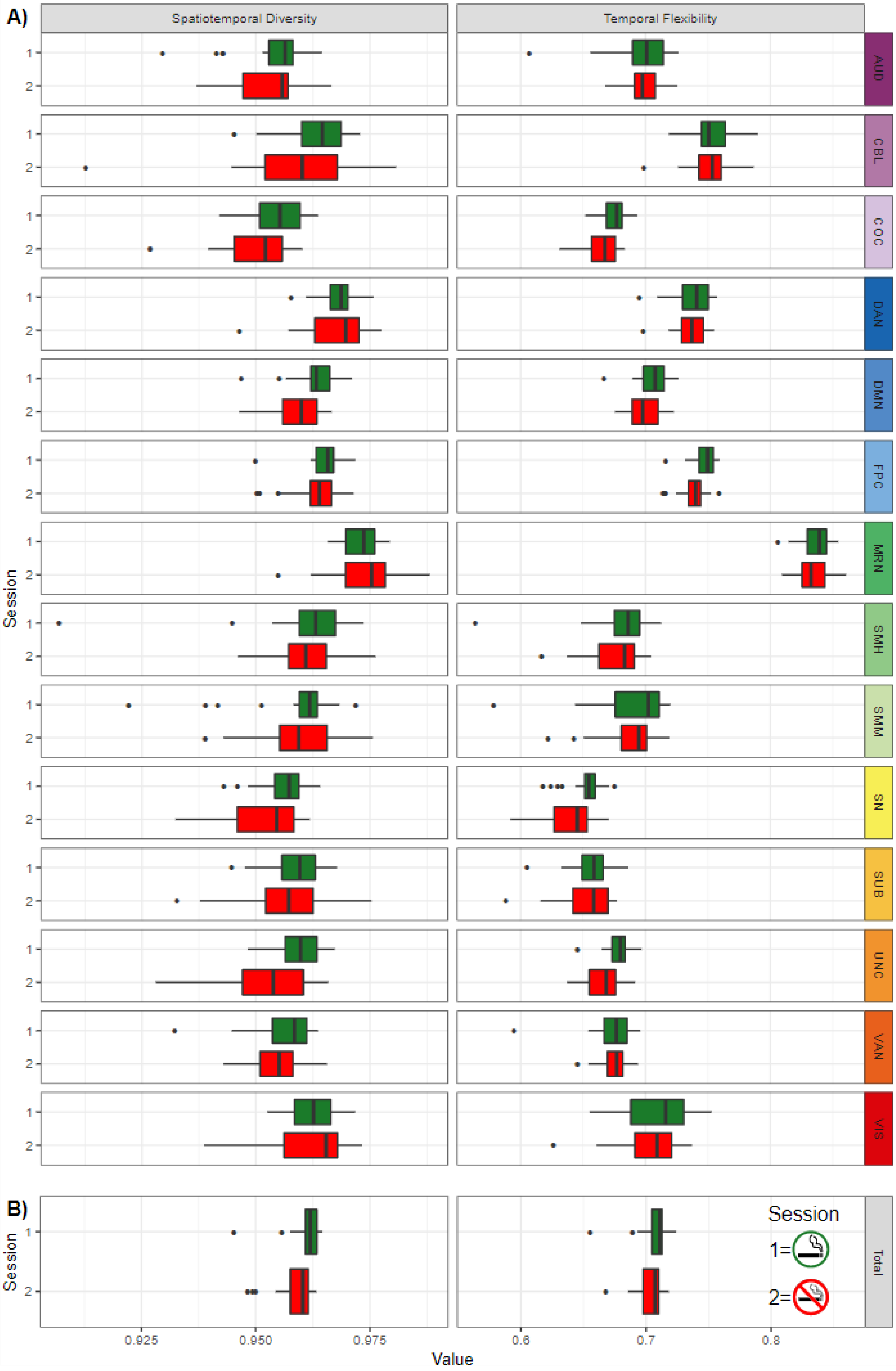

*Figure S4: Averaged time varying connectivity results A) averaged across nodes within each network and B) averaged across all networks. Green boxes=session 1 satiety; Red boxes=session 2 abstinence.*

*AUD=Auditory; CBL= Cerebellar; COC=Cingulo-Opercular Control; DAN=Dorsal Attention; DMN=Default Mode; FPC=Fronto-Parietal Control; MRN=Memory Retrieval; SMH=Somatomotor Hand; SMM=Somatomotor Mouth; SN= Salience; SUB=Subcortical; UNC=Uncategorized; VAN=Ventral Attention; VIS=Visual*

Figure S5: Nodes within each network showing state effect on time varying connectivity

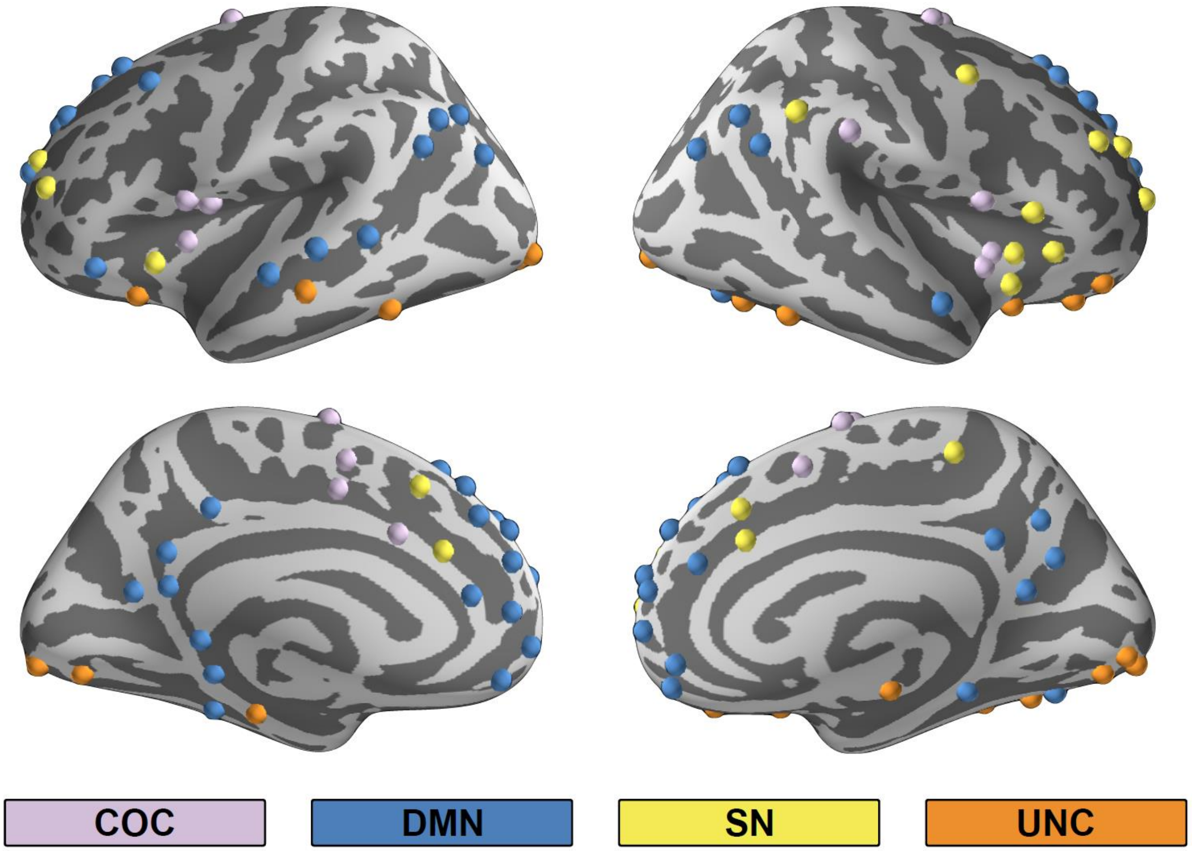


*Figure S5: Nodal membership for a priori networks showing STATE effect on STD*

*COC=Cingulo-Opercular Control; DMN=Default Mode; SN= Salience; UNC=Uncategorized*

Figure S6: *Time varying connectivity change as a function of abstinence displayed by group community segmentation*

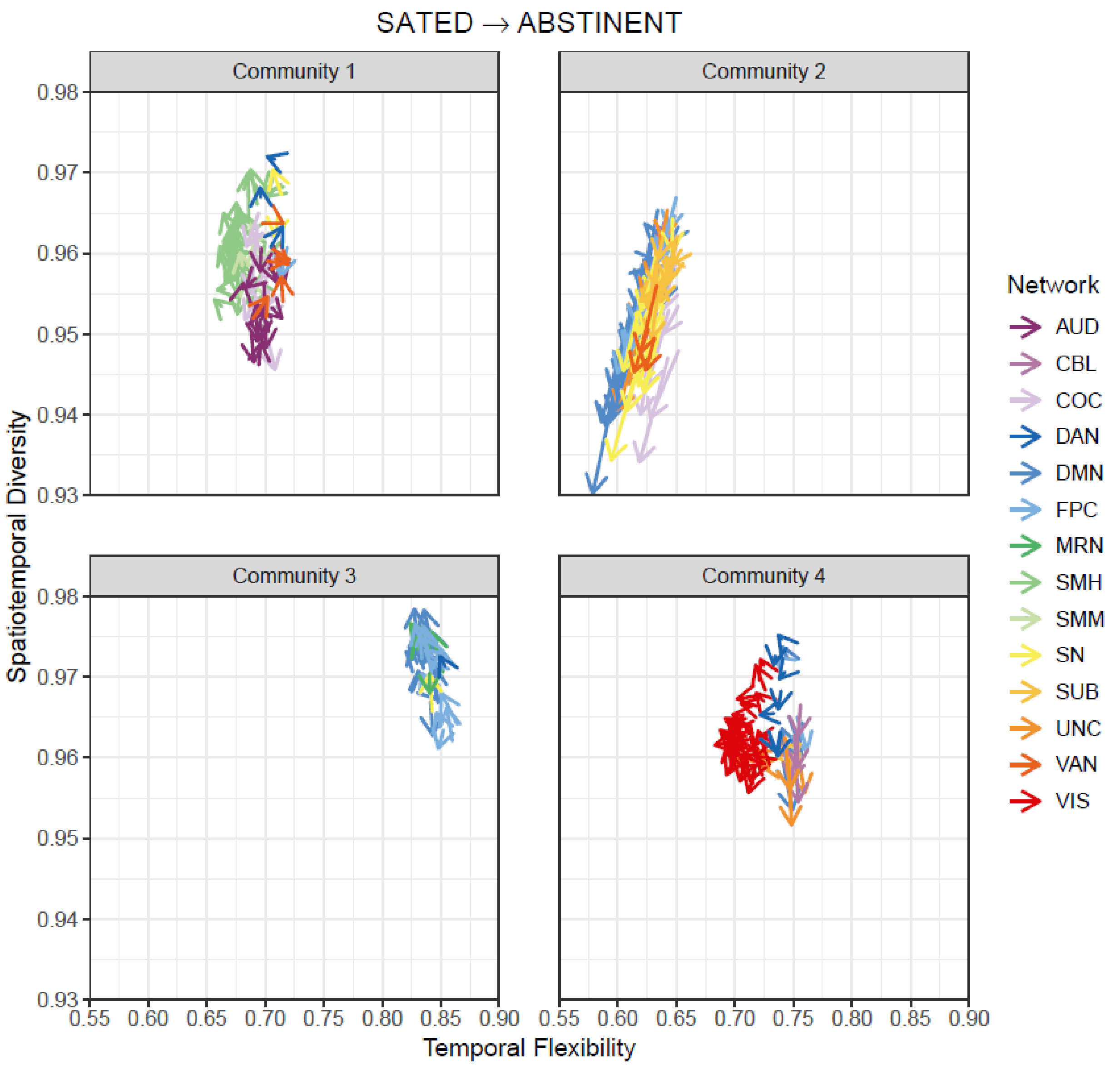


Figure S6: Color of vector indicates a priori network membership. Vectors indicate the magnitude of change in Temporal Flexibility (x-axis) and Spatiotemporal Diversity (y-axis) as a function of nicotine abstinence. Similar to a priori network results (Fig 4 manuscript), only one of the four communities shows a decrease in STD.

AUD=Auditory; CBL= Cerebellar; COC=Cingulo-Opercular Control; DAN=Dorsal Attention; DMN=Default Mode; FPC=Fronto-Parietal Control; MRN=Memory Retrieval; SMH=Somatomotor Hand; SMM=Somatomotor Mouth; SN= Salience; SUB=Subcortical; UNC=Uncategorized; VAN=Ventral Attention; VIS=Visual

Supplemental Tables:

Table S1:  240 Nodes in the Current Analysis

Table S2: 24 nodes excluded from current analysis

Table S1: 240 nodes included in the current analysis

| ROI | Network | Community | MNI x | MNI y | MNI z |
|---|---|---|---|---|---|
| 54 | AUD | 1 | 32 | -26 | 13 |
| 55 | AUD | 1 | 65 | -33 | 20 |
| 56 | AUD | 1 | 58 | -16 | 7 |
| 57 | AUD | 1 | -38 | -33 | 17 |
| 58 | AUD | 1 | -60 | -25 | 14 |
| 59 | AUD | 1 | -49 | -26 | 5 |
| 60 | AUD | 1 | 43 | -23 | 20 |
| 61 | AUD | 1 | -50 | -34 | 26 |
| 62 | AUD | 1 | -53 | -22 | 23 |
| 63 | AUD | 1 | -55 | -9 | 12 |
| 64 | AUD | 1 | 56 | -5 | 13 |
| 65 | AUD | 1 | 59 | -17 | 29 |
| 66 | AUD | 1 | -30 | -27 | 12 |
| 223 | CBL | 4 | -16 | -65 | -20 |
| 224 | CBL | 4 | -32 | -55 | -25 |
| 225 | CBL | 4 | 22 | -58 | -23 |
| 226 | CBL | 4 | 1 | -62 | -18 |
| 40 | COC | 1 | -3 | 2 | 53 |
| 41 | COC | 1 | 54 | -28 | 34 |
| 42 | COC | 1 | 19 | -8 | 64 |
| 43 | COC | 1 | -16 | -5 | 71 |
| 44 | COC | 1 | -10 | -2 | 42 |
| 45 | COC | 2 | 37 | 1 | -4 |
| 46 | COC | 1 | 13 | -1 | 70 |
| 47 | COC | 1 | 7 | 8 | 51 |
| 48 | COC | 1 | -45 | 0 | 9 |
| 49 | COC | 2 | 49 | 8 | -1 |
| 50 | COC | 2 | -34 | 3 | 4 |
| 51 | COC | 2 | -51 | 8 | -2 |
| 52 | COC | 2 | -5 | 18 | 34 |
| 53 | COC | 2 | 36 | 10 | 1 |
| 227 | DAN | 4 | 10 | -62 | 61 |
| 228 | DAN | 1 | -52 | -63 | 5 |
| 232 | DAN | 4 | 22 | -65 | 48 |
| 233 | DAN | 4 | 46 | -59 | 4 |
| 234 | DAN | 4 | 25 | -58 | 60 |
| 235 | DAN | 3 | -33 | -46 | 47 |
| 236 | DAN | 4 | -27 | -71 | 37 |
| 237 | DAN | 1 | -32 | -1 | 54 |
| 238 | DAN | 4 | -42 | -60 | -9 |
| 240 | DAN | 1 | 29 | -5 | 54 |
| 67 | DMN | 3 | -41 | -75 | 26 |
| 68 | DMN | 2 | 8 | 48 | -15 |
| 69 | DMN | 4 | -13 | -40 | 1 |
| 70 | DMN | 3 | -46 | -61 | 21 |
| 71 | DMN | 4 | 43 | -72 | 28 |
| 74 | DMN | 3 | -44 | -65 | 35 |
| 75 | DMN | 3 | -39 | -75 | 44 |

| | | | | | |
|---|---|---|---|---|---|
| 76 | DMN | 3 | -7 | -55 | 27 |
| 77 | DMN | 3 | 6 | -59 | 35 |
| 78 | DMN | 3 | -11 | -56 | 16 |
| 79 | DMN | 3 | -3 | -49 | 13 |
| 80 | DMN | 3 | 8 | -48 | 31 |
| 81 | DMN | 3 | 15 | -63 | 26 |
| 82 | DMN | 3 | -2 | -37 | 44 |
| 83 | DMN | 3 | 11 | -54 | 17 |
| 84 | DMN | 3 | 52 | -59 | 36 |
| 85 | DMN | 2 | 23 | 33 | 48 |
| 86 | DMN | 2 | -10 | 39 | 52 |
| 87 | DMN | 2 | -16 | 29 | 53 |
| 88 | DMN | 3 | -35 | 20 | 51 |
| 89 | DMN | 2 | 22 | 39 | 39 |
| 90 | DMN | 2 | 13 | 55 | 38 |
| 91 | DMN | 2 | -10 | 55 | 39 |
| 92 | DMN | 2 | -20 | 45 | 39 |
| 93 | DMN | 2 | 6 | 54 | 16 |
| 94 | DMN | 2 | 6 | 64 | 22 |
| 95 | DMN | 2 | -7 | 51 | -1 |
| 96 | DMN | 2 | 9 | 54 | 3 |
| 97 | DMN | 2 | -3 | 44 | -9 |
| 98 | DMN | 2 | 8 | 42 | -5 |
| 99 | DMN | 2 | -11 | 45 | 8 |
| 100 | DMN | 2 | -2 | 38 | 36 |
| 101 | DMN | 2 | -3 | 42 | 16 |
| 102 | DMN | 2 | -20 | 64 | 19 |
| 103 | DMN | 2 | -8 | 48 | 23 |
| 104 | DMN | 2 | -56 | -13 | -10 |
| 105 | DMN | 2 | -58 | -30 | -4 |
| 106 | DMN | 2 | 13 | 30 | 59 |
| 107 | DMN | 2 | 12 | 36 | 20 |
| 108 | DMN | 2 | 52 | -2 | -16 |
| 109 | DMN | 4 | -26 | -40 | -8 |
| 110 | DMN | 4 | 27 | -37 | -13 |
| 111 | DMN | 4 | -34 | -38 | -16 |
| 112 | DMN | 2 | 28 | -77 | -32 |
| 113 | DMN | 3 | 47 | -50 | 29 |
| 114 | DMN | 2 | -49 | -42 | 1 |
| 120 | DMN | 2 | -46 | 31 | -13 |
| 156 | FPC | 3 | -44 | 2 | 46 |
| 157 | FPC | 2 | 48 | 25 | 27 |
| 158 | FPC | 1 | -47 | 11 | 23 |
| 159 | FPC | 3 | -53 | -49 | 43 |
| 160 | FPC | 3 | -23 | 11 | 64 |
| 161 | FPC | 2 | 58 | -53 | -14 |
| 162 | FPC | 2 | 24 | 45 | -15 |
| 166 | FPC | 4 | 47 | 10 | 33 |
| 167 | FPC | 3 | -41 | 6 | 33 |
| 168 | FPC | 2 | -42 | 38 | 21 |
| 169 | FPC | 2 | 38 | 43 | 15 |
| 170 | FPC | 3 | 49 | -42 | 45 |
| 171 | FPC | 4 | -28 | -58 | 48 |

| | | | | | |
|---|---|---|---|---|---|
| 172 | FPC | 3 | 44 | -53 | 47 |
| 173 | FPC | 3 | 32 | 14 | 56 |
| 174 | FPC | 3 | 37 | -65 | 40 |
| 175 | FPC | 3 | -42 | -55 | 45 |
| 176 | FPC | 3 | 40 | 18 | 40 |
| 177 | FPC | 2 | -34 | 55 | 4 |
| 178 | FPC | 2 | -42 | 45 | -2 |
| 179 | FPC | 3 | 33 | -53 | 44 |
| 180 | FPC | 2 | 43 | 49 | -2 |
| 181 | FPC | 2 | -42 | 25 | 30 |
| 182 | FPC | 2 | -3 | 26 | 44 |
| 116 | MRN | 3 | -2 | -35 | 31 |
| 117 | MRN | 3 | -7 | -71 | 42 |
| 118 | MRN | 3 | 11 | -66 | 42 |
| 119 | MRN | 3 | 4 | -48 | 51 |
| 201 | MRN | 3 | 2 | -24 | 30 |
| 6 | SMH | 1 | -7 | -52 | 61 |
| 7 | SMH | 1 | -14 | -18 | 40 |
| 8 | SMH | 1 | 0 | -15 | 47 |
| 9 | SMH | 1 | 10 | -2 | 45 |
| 10 | SMH | 1 | -7 | -21 | 65 |
| 11 | SMH | 1 | -7 | -33 | 72 |
| 12 | SMH | 1 | 13 | -33 | 75 |
| 13 | SMH | 1 | -54 | -23 | 43 |
| 14 | SMH | 1 | 29 | -17 | 71 |
| 15 | SMH | 1 | 10 | -46 | 73 |
| 16 | SMH | 1 | -23 | -30 | 72 |
| 17 | SMH | 1 | -40 | -19 | 54 |
| 18 | SMH | 1 | 29 | -39 | 59 |
| 19 | SMH | 1 | 50 | -20 | 42 |
| 20 | SMH | 1 | -38 | -27 | 69 |
| 21 | SMH | 1 | 20 | -29 | 60 |
| 22 | SMH | 1 | 44 | -8 | 57 |
| 23 | SMH | 1 | -29 | -43 | 61 |
| 24 | SMH | 1 | 10 | -17 | 74 |
| 25 | SMH | 1 | 22 | -42 | 69 |
| 26 | SMH | 1 | -45 | -32 | 47 |
| 27 | SMH | 1 | -21 | -31 | 61 |
| 28 | SMH | 1 | -13 | -17 | 75 |
| 29 | SMH | 1 | 42 | -20 | 55 |
| 30 | SMH | 1 | -38 | -15 | 69 |
| 31 | SMH | 1 | -16 | -46 | 73 |
| 32 | SMH | 1 | 2 | -28 | 60 |
| 33 | SMH | 1 | 3 | -17 | 58 |
| 34 | SMH | 1 | 38 | -17 | 45 |
| 231 | SMH | 1 | 47 | -30 | 49 |
| 35 | SMM | 1 | -49 | -11 | 35 |
| 36 | SMM | 1 | 36 | -9 | 14 |
| 37 | SMM | 1 | 51 | -6 | 32 |
| 38 | SMM | 1 | -53 | -10 | 24 |
| 39 | SMM | 1 | 66 | -8 | 25 |
| 183 | SN | 1 | 11 | -39 | 50 |

| | | | | | |
|---|---|---|---|---|---|
| 184 | SN | 3 | 55 | -45 | 37 |
| 185 | SN | 1 | 42 | 0 | 47 |
| 186 | SN | 2 | 31 | 33 | 26 |
| 187 | SN | 2 | 48 | 22 | 10 |
| 188 | SN | 2 | -35 | 20 | 0 |
| 189 | SN | 2 | 36 | 22 | 3 |
| 190 | SN | 2 | 37 | 32 | -2 |
| 191 | SN | 2 | 34 | 16 | -8 |
| 192 | SN | 2 | -11 | 26 | 25 |
| 193 | SN | 2 | -1 | 15 | 44 |
| 194 | SN | 2 | -28 | 52 | 21 |
| 195 | SN | 2 | 0 | 30 | 27 |
| 196 | SN | 2 | 5 | 23 | 37 |
| 197 | SN | 2 | 10 | 22 | 27 |
| 198 | SN | 2 | 31 | 56 | 14 |
| 199 | SN | 2 | 26 | 50 | 27 |
| 200 | SN | 2 | -39 | 51 | 17 |
| 202 | SUB | 4 | 6 | -24 | 0 |
| 203 | SUB | 2 | -2 | -13 | 12 |
| 204 | SUB | 2 | -10 | -18 | 7 |
| 205 | SUB | 2 | 12 | -17 | 8 |
| 206 | SUB | 4 | -5 | -28 | -4 |
| 207 | SUB | 2 | -22 | 7 | -5 |
| 208 | SUB | 2 | -15 | 4 | 8 |
| 209 | SUB | 2 | 31 | -14 | 2 |
| 210 | SUB | 2 | 23 | 10 | 1 |
| 211 | SUB | 2 | 29 | 1 | 4 |
| 212 | SUB | 2 | -31 | -11 | 0 |
| 213 | SUB | 2 | 15 | 5 | 7 |
| 214 | SUB | 2 | 9 | -4 | 6 |
| 1 | UNC | 4 | -25 | -98 | -12 |
| 2 | UNC | 2 | 24 | 32 | -18 |
| 3 | UNC | 2 | -21 | -22 | -20 |
| 4 | UNC | 4 | 17 | -28 | -17 |
| 5 | UNC | 2 | 34 | 38 | -12 |
| 72 | UNC | 2 | -58 | -26 | -15 |
| 73 | UNC | 2 | 27 | 16 | -17 |
| 115 | UNC | 2 | -31 | 19 | -19 |
| 122 | UNC | 4 | 8 | -91 | -7 |
| 123 | UNC | 4 | 17 | -91 | -14 |
| 124 | UNC | 4 | -12 | -95 | -13 |
| 163 | UNC | 2 | -18 | -76 | -24 |
| 164 | UNC | 2 | 17 | -80 | -34 |
| 165 | UNC | 2 | 35 | -67 | -34 |
| 229 | UNC | 4 | -47 | -51 | -21 |
| 230 | UNC | 4 | 46 | -47 | -17 |
| 121 | VAN | 2 | -10 | 11 | 67 |
| 215 | VAN | 1 | 54 | -43 | 22 |
| 216 | VAN | 1 | -56 | -50 | 10 |
| 217 | VAN | 1 | -55 | -40 | 14 |
| 218 | VAN | 1 | 52 | -33 | 8 |
| 219 | VAN | 2 | 51 | -29 | -4 |

| | | | | | |
|---|---|---|---|---|---|
| 220 | VAN | 1 | 56 | -46 | 11 |
| 221 | VAN | 2 | 53 | 33 | 1 |
| 222 | VAN | 2 | -49 | 25 | -1 |
| 125 | VIS | 4 | 18 | -47 | -10 |
| 126 | VIS | 4 | 40 | -72 | 14 |
| 127 | VIS | 4 | 8 | -72 | 11 |
| 128 | VIS | 4 | -8 | -81 | 7 |
| 129 | VIS | 4 | -28 | -79 | 19 |
| 130 | VIS | 4 | 20 | -66 | 2 |
| 131 | VIS | 4 | -24 | -91 | 19 |
| 132 | VIS | 4 | 27 | -59 | -9 |
| 133 | VIS | 4 | -15 | -72 | -8 |
| 134 | VIS | 4 | -18 | -68 | 5 |
| 135 | VIS | 4 | 43 | -78 | -12 |
| 136 | VIS | 4 | -47 | -76 | -10 |
| 137 | VIS | 4 | -14 | -91 | 31 |
| 138 | VIS | 4 | 15 | -87 | 37 |
| 139 | VIS | 4 | 29 | -77 | 25 |
| 140 | VIS | 4 | 20 | -86 | -2 |
| 141 | VIS | 4 | 15 | -77 | 31 |
| 142 | VIS | 4 | -16 | -52 | -1 |
| 143 | VIS | 4 | 42 | -66 | -8 |
| 144 | VIS | 4 | 24 | -87 | 24 |
| 145 | VIS | 4 | 6 | -72 | 24 |
| 146 | VIS | 4 | -42 | -74 | 0 |
| 147 | VIS | 4 | 26 | -79 | -16 |
| 148 | VIS | 4 | -16 | -77 | 34 |
| 149 | VIS | 4 | -3 | -81 | 21 |
| 150 | VIS | 4 | -40 | -88 | -6 |
| 151 | VIS | 4 | 37 | -84 | 13 |
| 152 | VIS | 4 | 6 | -81 | 6 |
| 153 | VIS | 4 | -26 | -90 | 3 |
| 154 | VIS | 4 | -33 | -79 | -13 |
| 155 | VIS | 4 | 37 | -81 | 1 |

ROI= ROI number from Power et al., 2011 parcellation; Community= arbitrary community membership based on group community structure

*Network abbreviations: AUD=Auditory; CBL= Cerebellar; COC=Cingulo-Opercular Control; DAN=Dorsal Attention; DMN=Default Mode; FPC=Fronto-Parietal Control; MRN=Memory Retrieval; SMH=Somatomotor Hand; SMM=Somatomotor Mouth; SN= Salience; SUB=Subcortical; UNC=Uncategorized; VAN=Ventral Attention; VIS=Visual*

Table S2: 24 nodes excluded from current analysis

| ROI | Network | MNI x | MNI y | MNI z |
|---|---|---|---|---|
| 75 | DMN | 6 | 67 | -4 |
| 78 | DMN | -18 | 63 | -9 |
| 81 | DMN | -44 | 12 | -34 |
| 82 | DMN | 46 | 16 | -30 |
| 83 | DMN | -68 | -23 | -16 |
| 116 | DMN | 65 | -12 | -19 |
| 119 | DMN | 65 | -31 | -9 |
| 120 | DMN | -68 | -41 | -5 |
| 128 | DMN | 52 | 7 | -30 |
| 129 | DMN | -53 | 3 | -27 |
| 139 | DMN | 49 | 35 | -12 |
| 181 | FPC | 34 | 54 | -13 |
| 2 | UNC | 27 | -97 | -13 |
| 4 | UNC | -56 | -45 | -24 |
| 5 | UNC | 8 | 41 | -24 |
| 8 | UNC | -37 | -29 | -26 |
| 9 | UNC | 65 | -24 | -19 |
| 10 | UNC | 52 | -34 | -27 |
| 11 | UNC | 55 | -31 | -17 |
| 182 | UNC | -21 | 41 | -20 |
| 247 | UNC | 33 | -12 | -34 |
| 248 | UNC | -31 | -10 | -36 |
| 249 | UNC | 49 | -3 | -38 |
| 250 | UNC | -50 | -7 | -39 |

ROI= ROI number from Power et al., 2011 parcellation

Network abbreviations: DMN=Default Mode; FPC=Fronto-Parietal Control; UNC=Uncategorized

*REFERENCES*


1. Esteban O, Markiewicz CJ, Blair RW, Moodie CA, Isik AI, Erramuzpe A, Kent JD, Goncalves M, DuPre E, Snyder M, Oya H, Ghosh SS, Wright J, Durnez J, Poldrack RA, Gorgolewski KJ. fMRIPrep: a robust preprocessing pipeline for functional MRI. Nat Meth. 2018; doi:10.1038/s41592-018-0235-4

2. Esteban, Oscar, Blair, Ross, Markiewicz, Christopher J., Berleant, Shoshana L., Moodie, Craig, Ma, Feilong, Isik, Ayse Ilkay, Erramuzpe, Asier, Kent, James D., Goncalves, Mathias, DuPre, Elizabeth, Sitek, Kevin R., Gomez, Daniel E. P., Lurie, Daniel J., Ye, Zhifang, Salo, Taylor, Valabregue, Romain, Amlien, Inge K., Liem, Franz, Jacoby, Nir, Stojic, Hrvoje, Halchenko, Yaroslav O., Rivera-Dompenciel, Adriana, Ciric, Rastko, Sneve, Markus H., Heinsfeld, Anibal S., Thompson, William H, Tooley, Ursula A., Poldrack, Russell A., Gorgolewski, Krzysztof J. fMRIPrep: a robust preprocessing pipeline for functional MRI. Zenodo; 2019. Available

from: https://doi.org/10.5281/zenodo.3511440.

3. Gorgolewski K, Burns CD, Madison C, Clark D, Halchenko YO, Waskom ML, Ghosh SS. Nipype: a flexible, lightweight and extensible neuroimaging data processing framework in python. Front Neuroinform. 2011 Aug 22;5(August):13. doi:10.3389/fninf.2011.00013.

4. Gorgolewski KJ, Esteban O, Ellis DG, Notter MP, Ziegler E, Johnson H, Hamalainen C, Yvernault B, Burns C, Manhães-Savio A, Jarecka D, Markiewicz CJ, Salo T, Clark D, Waskom M, Wong J, Modat M, Dewey BE, Clark MG, Dayan M, Loney F, Madison C, Gramfort A, Keshavan A, Berleant S, Pinsard B, Goncalves M, Clark D, Cipollini B, Varoquaux G, Wassermann D, Rokem A, Halchenko YO, Forbes J, Moloney B, Malone IB, Hanke M, Mordom D, Buchanan C, Pauli WM, Huntenburg JM, Horea C, Schwartz Y, Tungaraza R, Iqbal S, Kleesiek J, Sikka S, Frohlich C, Kent J, Perez-Guevara M, Watanabe A, Welch D, Cumba C, Ginsburg D, Eshaghi A, Kastman E, Bougacha S, Blair R, Acland B, Gillman A, Schaefer A, Nichols BN, Giavasis S, Erickson D, Correa C, Ghayoor A, Küttner R, Haselgrove C, Zhou D, Craddock RC, Haehn D, Lampe L, Millman J, Lai J, Renfro M, Liu S, Stadler J, Glatard T, Kahn AE, Kong X-Z, Triplett W, Park A, McDermottroe C, Hallquist M, Poldrack R, Perkins LN, Noel M, Gerhard S, Salvatore J, Mertz F, Broderick W, Inati S, Hinds O, Brett M, Durnez J, Tambini A, Rothmei S, Andberg SK, Cooper G, Marina A, Mattfeld A, Urchs S, Sharp P, Matsubara K, Geisler D, Cheung B, Floren A, Nickson T, Pannetier N, Weinstein A, Dubois M, Arias J, Tarbert C, Schlamp K, Jordan K, Liem F, Saase V, Harms R, Khanuja R, Podranski K, Flandin G, Papadopoulos Orfanos D, Schwabacher I, McNamee D, Falkiewicz M, Pellman J, Linkersdörfer J, Varada J, Pérez-García F, Davison A, Shachnev D, Ghosh S. Nipype: a flexible, lightweight and extensible neuroimaging data processing framework in Python. 2017. doi:10.5281/zenodo.581704.
